# VSS: Variance-stabilized signals for sequencing-based genomic signals

**DOI:** 10.1101/2020.01.31.929174

**Authors:** Faezeh Bayat, Maxwell Libbrecht

## Abstract

**Motivation:** A sequencing-based genomic assay such as ChIP-seq outputs a real-valued signal for each position in the genome that measures the strength of activity at that position. Most genomic signals lack the property of variance stabilization. That is, a difference between 100 and 200 reads usually has a very different statistical importance from a difference between 1,100 and 1,200 reads. A statistical model such as a negative binomial distribution can account for this pattern, but learning these models is computationally challenging. Therefore, many applications—including imputation and segmentation and genome annotation (SAGA)—instead use Gaussian models and use a transformation such as log or inverse hyperbolic sine (asinh) to stabilize variance.

**Results:** We show here that existing transformations do not fully stabilize variance in genomic data sets. To solve this issue, we propose VSS, a method that produces variance-stabilized signals for sequencing-based genomic signals. VSS learns the empirical relationship between the mean and variance of a given signal data set and produces transformed signals that normalize for this dependence. We show that VSS successfully stabilizes variance and that doing so improves downstream applications such as SAGA. VSS will eliminate the need for downstream methods to implement complex mean-variance relationship models, and will enable genomic signals to be easily understood by eye.

**Availability:** https://github.com/faezeh-bayat/Variance-stabilized-units-for-sequencing-based-genomic-signals.

## 1 Introduction

Sequencing-based assays can measure many types of genomic biochemical activity, including transcription factor binding, histone modifications and chromatin accessibility. These assays work by extracting DNA fragments from a sample that exhibit the desired type of activity, sequencing the fragments to produce sequencing reads and mapping each read to the genome. Each of these assays produces a genomic signal— that is, a signal that has a value for each base pair in the genome. Examples include ChIP-seq measurements of transcription factor binding or histone modification and measurements of chromatin accessibility from DNase-seq, FAIRE-seq or ATAC-seq. The natural unit of sequencing-based assays is the read count: the number of reads that mapped to a given position in the genome (after extending and shifting; see Methods).

Read counts of genomic assays have a nonuniform mean-variance relationship, which poses a challenge to their analysis. For example, a locus with 1,000 reads in one experiment might get 1,100 reads in a replicate experiment by chance, whereas a locus with 100 reads might usually see no more than 110 reads by chance in a replicate. This property means that, for example, the difference in read count between biosamples is a poor measure of the difference in activity. To handle this issue, most statistical models of genomic signals— such as those used in peak calling—model the mean-variance relationship of read counts explicitly using, for example, a negative binomial distribution [1–11].

However, negative binomial models are challenging to implement and optimize, so many methods resort to Gaussian models. Two prominent examples include segmentation and genome annotation (SAGA) methods, such as Segway or IDEAS [12–16], and imputation methods such as ChromImpute, PREDICTD and Avocado [17–19]. In the former example, many SAGA methods use a Gaussian distribution to model the distribution of genomic signals given a certain annotation label (others binarize signal [20] or use a negative binomial read count model [21]). In the latter example, imputation methods optimize a mean squared error (MSE) objective function, which is equivalent to log likelihood in a Gaussian model.

Most Gaussian-based methods employ a variance-stabilizing transformation to handle the nonuniform mean-variance relationship. They most commonly use the log or inverse hyperbolic sine transformations (asinh), which have the formulae log(*x* + *c*) for a constant *c* (usually 1) and 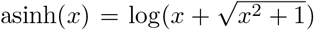 respectively [22].

Variance-stabilizing transformations are also required for visualizing genomic signals. As previously noted, Euclidean distances in a 2D plot correspond to log likelihood in a Gaussian model, so a nonuniform meanvariance relationship complicates visualization. Therefore, most visualizations of genomic signals such as genome browsers [23] employ a variance-stabilizing transformation such as log.

Despite the widespread use of log and asinh transformations to stabilize variance, to our knowledge, no work has evaluated whether they in fact do so. The use of these transformations assumes that the signals have a specific mean-variance relationship (Methods). Here we show that, for many genomic signals, this assumptions is violated and thus existing transformations do not fully stabilize variance (Results). To solve this issue, we present VSS, a method that produces variance-stabilized genomic signals. VSS determines the empirical mean-variance relationship of a genomic signal by comparing replicates. It uses this empirical mean-variance relationship to produce a transformation function that precisely stabilizes variance.

VSS source code is available at https://github.com/faezeh-bayat/Variance-stabilized-units-for-sequencing-based-genomic-signals.

### 1.1 Related work

Three methods have been developed to normalize sequencing-based genomic signals across experiments. First, fold enrichment measures a genomic signal as the ratio of reads of the experiment to a control (such as ChIP Input) [13]. Second, Poisson p-value measures a signal as the log p-value of a Poisson distribution test with a null hypothesis derived from a control distribution [24]. Third, S3norm [25] normalizes a collection of data sets by matching their empirical sequencing depth and signal-noise ratio. However, these methods normalize the mean of signals but do not stabilize variance (Results).

Many of the challenges mentioned here also exist for assays of gene expression such as RNA-seq data [22, 26, 27, 2, 28–31, 6, 32–34]. In particular, the method voom [31] uses the empirical mean-variance to stabilize variance of RNA-seq, like VSS. However, voom does not apply to genomic signals such as ChIP-seq and ATAC-seq.

## 2 Methods

### 2.1 ChIP-seq data

We acquired ChIP-seq data from the ENCODE consortium (encodeproject.org, Supplementary Section A) for the histone modification H3K4me3 on eleven cell lines: GM12878, H1-hESC, HUVEC, K562, NHLF, GM06990, HCPEpiC, AG09319, NHEK, HMEC and HSMM. We also used histone modifications H3K36me3, H3K4me1, H3K27me3 and H3K9me3 on H1-hESC cell line. Histone modification H2AFZ were used on cell lines NHEK and HSMM. In addition, we used histone modification H3K79me2 on NHEK, HSMM and HMEC cell lines. We also used histone modification H3K9me3 on four cell lines: NHEK, AG04450, HMEC and HSMM. Finally, H3K36me3 histone modification were used on HMEC cell line. ENCODE accession number of these assays are provided in the Supplementary Information Table 1. These ChIP-seq data sets were processed with a uniform pipeline [35]. Briefly, the ChIP-seq reads were mapped to the hg19 reference genome and reads were shifted and extended according to the estimated fragment length to produce a read count for each genomic position. As controls, ChIP-seq Input experiments were performed by the same labs. Two signals were produced: fold enrichment and log p-value. Fold enrichment signal is defined as the ratio of observed data over control [13]. P-value signal is defined as the log p-value of a Poisson model with a null distribution derived from the control [24].

### 2.2 RNA-seq data

For use in evaluation, we acquired RNA-seq data sets for each of the cell types above from the Roadmap Epigenomics consortium [24]. These RNA-seq data sets were processed with a uniform pipeline that produces a TPM value for each gene [24]. To stabilize the variance of these signals, we used an asinh transformation.

### 2.3 Identifying the mean-variance relationship

Our variance-stabilizing transformation depends on determining the mean-variance relationship for the input data set. We learn this relationship by comparing multiple replicates of the same experiment. We designate two replicates as the *base* and *auxiliary* replicates, respectively. The following process is iterated for all possible choices of base and auxiliary (see below).

Let the observed signal at position *i* be 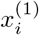 and 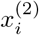 for the base and auxiliary replicates respectively. Our model imagines that every position *i* has an unknown distribution of sequencing reads for the given assay *x_i_*, which has mean *μ_i_* = mean(*x_i_*). We further suppose that there is a relationship *σ*(*μ*) between the mean and variance of these distributions. That is, var(*x_i_*) = *σ*(*μ_i_*)^2^. We are interested in learning *σ*(*μ*). Observe that *x_i_* is an unbiased estimate of *μ_i_*, and that 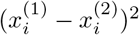 is an unbiased estimate of *σ*(*μ_i_*)^2^. We use this observation to estimate the function *σ*(*μ*) as follows.

We first sort the *N* genomic positions *i* ∈ {1…*N*} by the value of 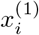 and define bins with *b* genomic positions each. Let *I_j_* ⊆ {1…*N*} be the set of positions in bin *j.* For each bin *j*, we compute 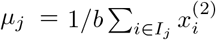 and 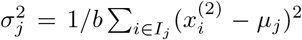. To increase the robustness of these estimates, we smooth across bins by defining

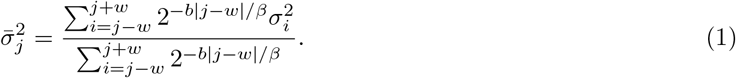

That is, we take the weighted average of 2*w* + 1 bins centered on *j*, where bin *j* + *k* has weight 2^-*bk/β*^. *β* is a bandwidth parameter—a high value of *β* means that weight is spread over many bins, whereas a low value means that weight in concentrated on a small number of bins. We define the window size w such that it includes bins with weight at least 0.01; specifically, *w* = –*β*log(0.01)/*b* log(2).

The choice of *b* and *β* forms a bias-variance trade-off. Larger values of *b* and *β* lead to more observations contributing to each estimate 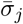 and therefore result in a lower variance. In contrast, small values of *b* and *β* lead to a very homogeneous set of positions *I_j_* and therefore less averaging across dissimilar positions.

Most genomic signals are zero-inflated. That is, a large fraction of positions have zero signal. To account for this pattern, we defined a separate bin for zero-signal positions 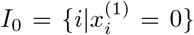 and defined *σ*_0_ and *μ*_0_ as above. We used this zero bin for raw and fold enrichment (FE) signals, but not log Poisson p-value (LPPV), which are not zero-inflated.

We used a smoothing spline to fit an estimated mean-variance curve 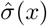. A smoothing spline estimator implements a regularized regression over the natural spline basis. We fit a function 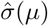 using the estimated values of 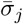. The spline coefficients *w* are selected to minimize

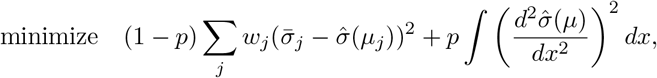

where *μ* and 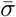 are a set of observations obtained from mean-variance data points. Variables 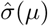, *w*, *p* represent smooth spline curve, weight coefficients and smoothing parameter respectively. The variable *p* parameter varies between (0,1] such that *p* = 0 results in a cubic spline with no smoothing, and when *p* approaches zero the result is a linear function. To find the optimum value of *spar* parameter (*p*), first the smooth.spline function is called by activating the cross-validation in the smooth.spline (CV=TRUE). Following the cross-validation procedure, *spar* parameter is returned as the smoothing factor. We identified the optimal curve using the R function call `smooth.spline(means, sigmas, spar=p`).

The process above assumed a fixed choice of base and auxiliary replicates. We repeat this process for each possible choice of base and auxiliary replicates (that is, for *M* replicates, the process above is repeated *M*(*M* – 1) times) and aggregate sufficient statistics across all iterations to produce an aggregated 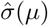.

We performed a hyperparameter search to choose *b* and *β* (Supplementary Figure 7 and Figure 8), using the log likelihood and variance instability metrics (defined below). We chose *β* = 10^3^ and *b* = 10^5^ for zero-inflated signals (raw and fold enrichment) and *β* = 10^7^ and *b* = 10^3^ for non-zero-inflated signals (log Poisson p-value).

### 2.4 Calculating variance-stabilized signals

Having learned the mean-variance relationship, we compute VSS using the variance-stabilizing transformation [36]

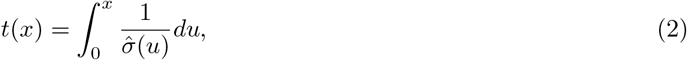

where *x* is an untransformed signal and 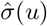 is the learned standard deviation for a signal with mean *u*. This transformation is guaranteed to be variance-stabilizing; that is, var(*t*(*x_i_*)) is constant for all genomic positions *i*.

### 2.5 Alternative transformations

To attempt to stabilize the variance, existing methods usually apply either a log or arcsinh transformation. These transformations are used because they are variance-stabilizing for certain mean-variance relationships [37]. Specifically, log(*x*) is variance-stabilizing when *σ*(*μ*) = *sμ* for some constant *s*, and arcsinh(*x*) is variance-stabilizing when 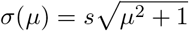 [38]. In the experiments below, we compare to both existing units under both existing transformations. We evaluated the performance log with a general linear transformation (log(*ax* + *b*)). We found that doing so did not improve results (Supplementary Section C), so we focused on the standard offset log(*x* +1).

### 2.6 Variance quality-of-fit evaluation

A transformation implicitly assumes that a data set has a specific mean-variance relationship. The assumed variance 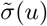 for a given value u equals the inverse of the derivative of the transformation (Section 2.4)

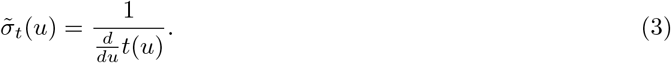

As noted above, a log(*x* + 1) transformation implicitly assumes the mean-variance relationship 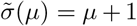 and the arcsinh(*x*) transformation assumes the mean-variance relationship 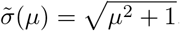.

To measure the quality of fit of an assumed mean-variance relationship, we evaluated the data log likelihood under the assumed 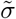. Specifically, the log likelihood of a given data set is defined as

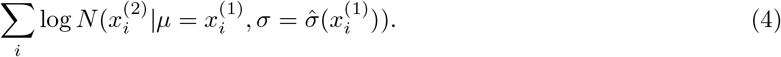

A Gaussian distribution appears in this expression because that is the max-entropy distribution with a specific mean and variance. This value is maximized when the inferred variance equals the variance of the data.

### 2.7 Variance instability evaluation

To evaluate whether a given transformation achieves a uniform mean-variance relationship, we defined the following variance instability metric. Let 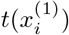 and 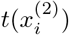 be the transformed signals at the *i*th genomic position. Using the binning approach described above, we divided genomic positions to *B* bins of increasing value of 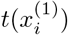, where each bin is of size *b* = 10000.

Let *v_j_* be the mean squared difference between replicates for positions in bin *j*,

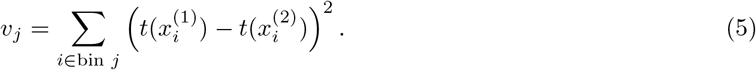

Let *σ*_1_ and *σ*_2_ be the standard deviation of *t*(*x*^1^) and *t*(*x*^2^) respectively. We define the variance instability metric as the scaled variance of *v_j_* across bins,

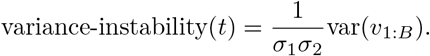

The 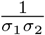 factor normalizes for the variance of the transformed signal; without this factor, *t*(*x*) and *αt*(*x*) (for a constant *α*) have different variance instability. Signals with unstable variance will have large values of the variance instability metric.

### 2.8 Segmentation and genome annotation (SAGA) evaluation

As described above, segmentation and genome annotation (SAGA) algorithms are sensitive to the meanvariance relationship in the input data sets. SAGA algorithms take as input a collection of signals for a given biosample. They partition the genome and assign a label to each segment such that positions with the same label have similar patterns in the input data sets. SAGA algorithms are widely used to integrate data sets and annotate regulatory elements.

To evaluate the quality of annotations produced by signals under a given transformation, we defined the following SAGA metric. Following previous work [16] we quantified quality of a SAGA annotation according to the strength of the relationship between the annotation of a genic region with that gene’s expression. Specifically, for a collection of signals from a given biosample, we used the SAGA algorithm Segway [12] to produce an annotation. This annotation assigns one of k integer labels *l_i_* ∈ {1..*k*} to each genomic position *i*. We defined features for each gene as follows. We divided each genic region into 20 bins by dividing the transcribed region into 10 equally-spaced bins and defining five 1kb bins upstream of the transcription start site (TSS) and downstream of the transcription termination site (TTS) respectively. We defined features *f_b,k_* for each bin b as a one-hot encoding of the majority label in each bin. That is, this process associates each gene with a vector of 20*k* features.

We trained an Extreme Gradient Boosting (XGBoost) regression model to predict a gene’s RNA-seq expression (Section “Segmentation and genome annotation (SAGA) evaluation”) value from this vector of features. We trained a regression model on a matrix containing all genes in a chromosome. As features we used a one-hot encoding feature vector that indicates 1 in the corresponding position of the predicted label and 0 elsewhere. For each bin, we considered the majority feature vector as a representation of that bin’s annotation. We used the coefficient of determination (*r*^2^) to quantify the predictive power of this regressor.

For each transformation method, we used four different values *k* = {3,5,10,15} for number of the labels to be predicted by annotation. We considered four biosamples: H1-hESC, NHEK, HSMM and HMEC. We used all available replicated histone modification data for each biosample: We used H3K36me3, H3K4me3, H3K9me3, H3K27me3 and H3K4me1 for H1-hESC. We used H2AFZ, H3K4me3, H3K9me3 and H3K79me2 for NHEK. We used H2AFZ, H3K4me3, H3K9me3 and H3K79me2 for HSMM. We used H3K36me3, H3K4me3, H3K9me3 and H3K79me2 for HMEC.

## 3 Results

### 3.1 Genomic signals are not variance-stabilized

To evaluate whether existing units for genomic signals have stable variance, we computed the mean-variance relationship for a number of existing data sets (Figure 1c). As we expected, we found that the variance has a strong dependence on the mean; genomic positions with low signals experience little variance across replicates, whereas positions with high signals experience much larger variance (Figure 1c). Moreover, the relationship does not match that expected by the currently-used log(*x* + 1) and asinh(*x*) transformations. A transformation implicitly imposes a specific mean-variance relationship; the inferred variance for a given value equals the inverse of the derivative of the transformation (Methods). For example, the former transformation assumes a linear relationship (Methods). The observed mean-variance relationship does not precisely match the relationships assumed by either transformation, indicating that neither of these transformations is fully variance-stabilizing (Figure 1d).

**Fig. 1.**
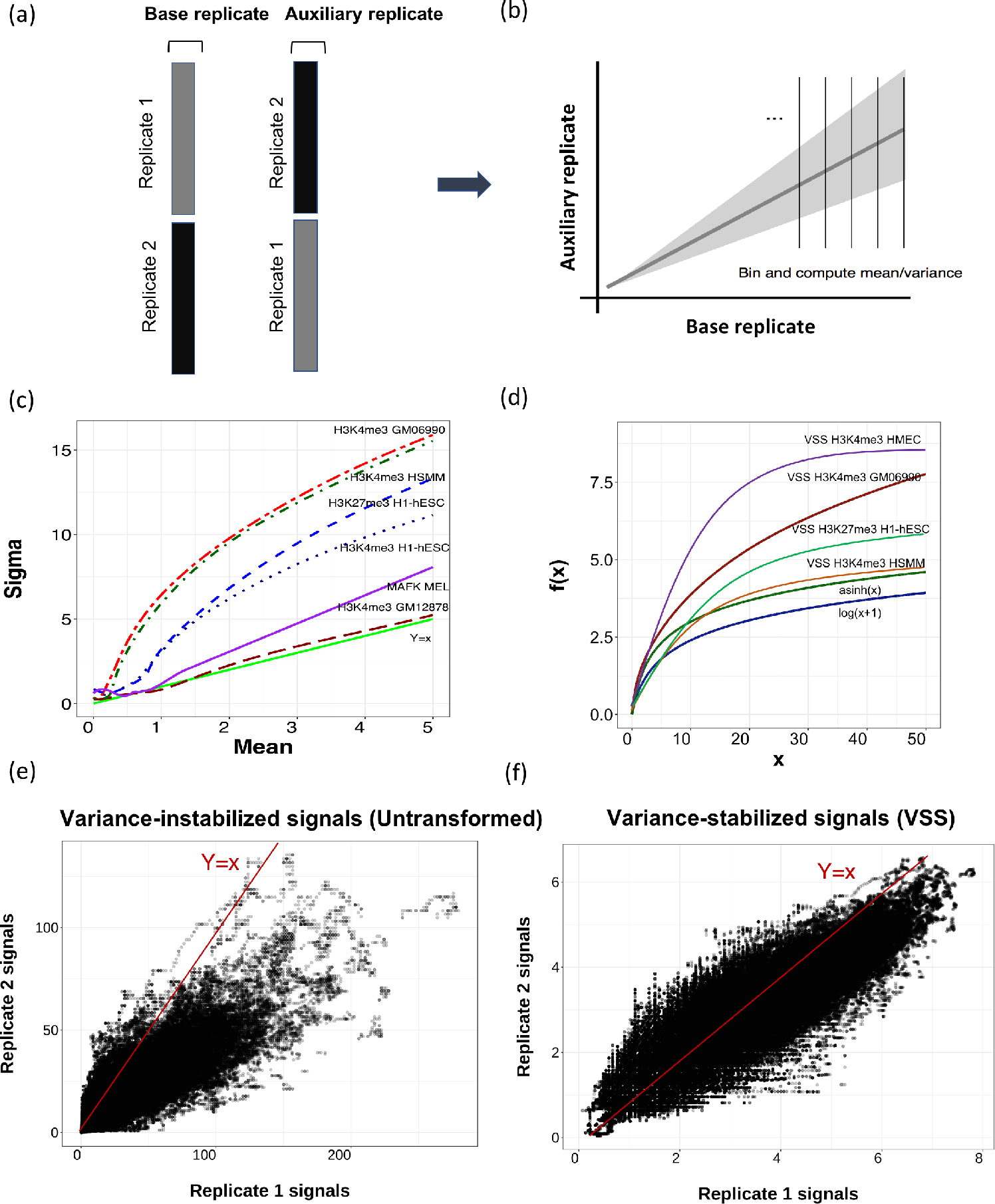
General schematic of the VSS method. (a,b) VSS uses two or more replicates to learn the empirical mean-variance relationship of the input data. It does so by binning values according the *base* replicate and evaluating the variance in the *auxiliary* replicate. (c) Learned mean-variance relationships for several data sets. Horizontal and vertical axes denote mean and standard deviation respectively. Note that the mean-variance relationship differs across data sets, indicating that each requires a different transformation. (d) Learned transformation functions. Horizontal and vertical axes indicate input and output values respectively. (e,f) Replicate 1 vs Replicate 2 signals in H3K4me3 HSMM before (e) and after (f) VSS transformation.

The observation that existing transformations are not variance-stabilizing was confirmed when we quantified this fit (Figure 2). A transformation implicitly assumes that a data set has a specific mean-variance relationship; for example, a log transform assumes a linear mean-variance relationship. To measure the accuracy of a variance estimate, we used the log likelihood of a given mean-variance relationship estimate, which is maximized when the inferred variance equals the variance of the data (Methods). As expected, we found that a uniform variance model implied by using untransformed signals had a poor likelihood (average log density of ™1.9), reflecting non-uniform variance (Figure 2b). We found that the variance estimates from the log(*FE*+1) and asinh(FE), where FE is the Fold enrichment signal, greatly improved the likelihood (average log density of −1.3 and −1.5 respectively). However, we found that mean-variance relationship learned by VSS had much better likelihood (average log density −1.2) than either transformation, indicating that the learned curve successfully models the mean-variance relationship of the data (Figure 2b). We found that VSS’s mean-variance fit was also better than log or asinh when using either raw reads or log Poisson p-value (LPPV) as the base units (Figure 2c,d).

**Fig. 2.**
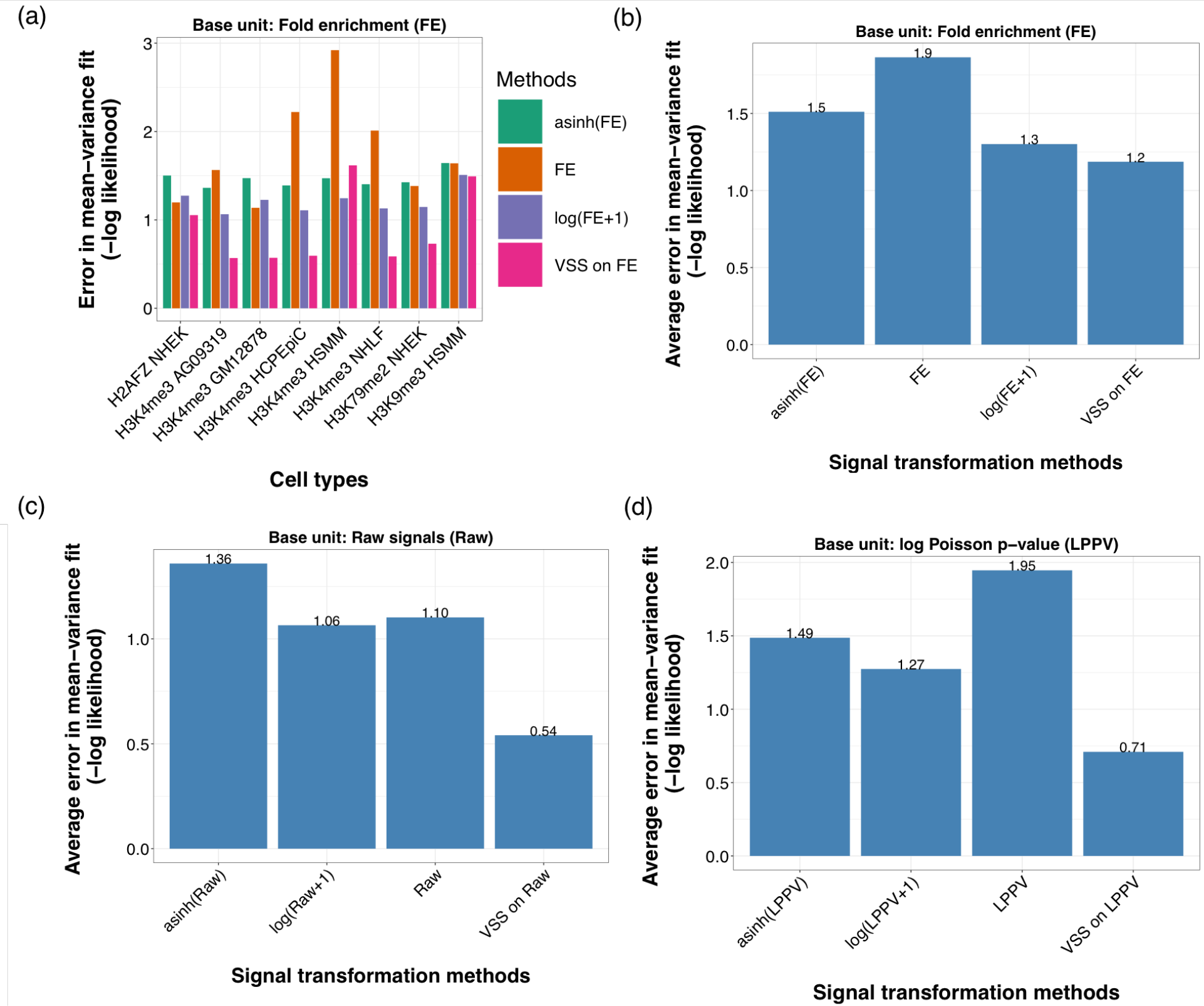
(a) Goodness of fit to the mean-variance relationship, measured by Gaussian log likelihood (Methods). Lower values of negative log likelihood indicates better fit. Log likelihood was computed on chromosome 21; VSS’s meanvariance relationship was trained on chromosome 22. (b) Same as (a), but averaged across data sets.

Moreover, we found that the mean-variance relationship differs greatly between experiments. For many histone modification ChIP-seq experiments like H3K4me3 HSMM and H3K9me3 HUVEC, a log transformation yields nearly-optimal fit, indicating that the data has a nearly linear mean-variance relationship (Figure 2a). However, other experiments like H3K4me3 in GM12878 and H2AFZ in NHEK, have a very non-linear mean-variance relationship (Figure 2a). In fact, for some experiments, a log or asinh transformation has worse fit than no transformation, indicating that these transformations actually destabilize the variance (Figure 2a). Future work should investigate what properties of an experiment determine its mean-variance relationship. The mean-variance relationship learned by VSS correctly captures these differences, as indicated by its good likelihood on all data sets. These differences indicates that it is necessary to learn a separate mean-variance relationship for each data set, rather than applying a single transformation (such as log or asinh) to every data set.

### 3.2 Differences between replicates are stabilized after transformation

To measure whether a given transformation stabilizes variance in a given signal data set, we defined the variance-instability metric (Methods). This metric is defined as the variance of mean squared between- replicate differences across bins defined by signal value. Signals with unstable variance will have a large value for this metric.

We found that signals transformed using VSS have better variance stability than either untransformed signals or signals after alternative transformations (Figure 3). Fold enrichment (FE) signals transformed by either log(*x* + 1) and asinh(*x*) on had an average of 1.8 variance instability, whereas VSS have instability of 1.5. Changing the offset of the log transformation—log(*ax* + *b*)—did not substantially improve results for any choice of *a* or *b* (Supplementary Information 9 and 10). This indicates that VSS units have more consistent signals among different replicates of an experiment (Figure 3d and Figure 3e). This pattern also holds when using Raw or LPPV as the base signal (Figure 3f and Figure 3g).

**Fig. 3.**
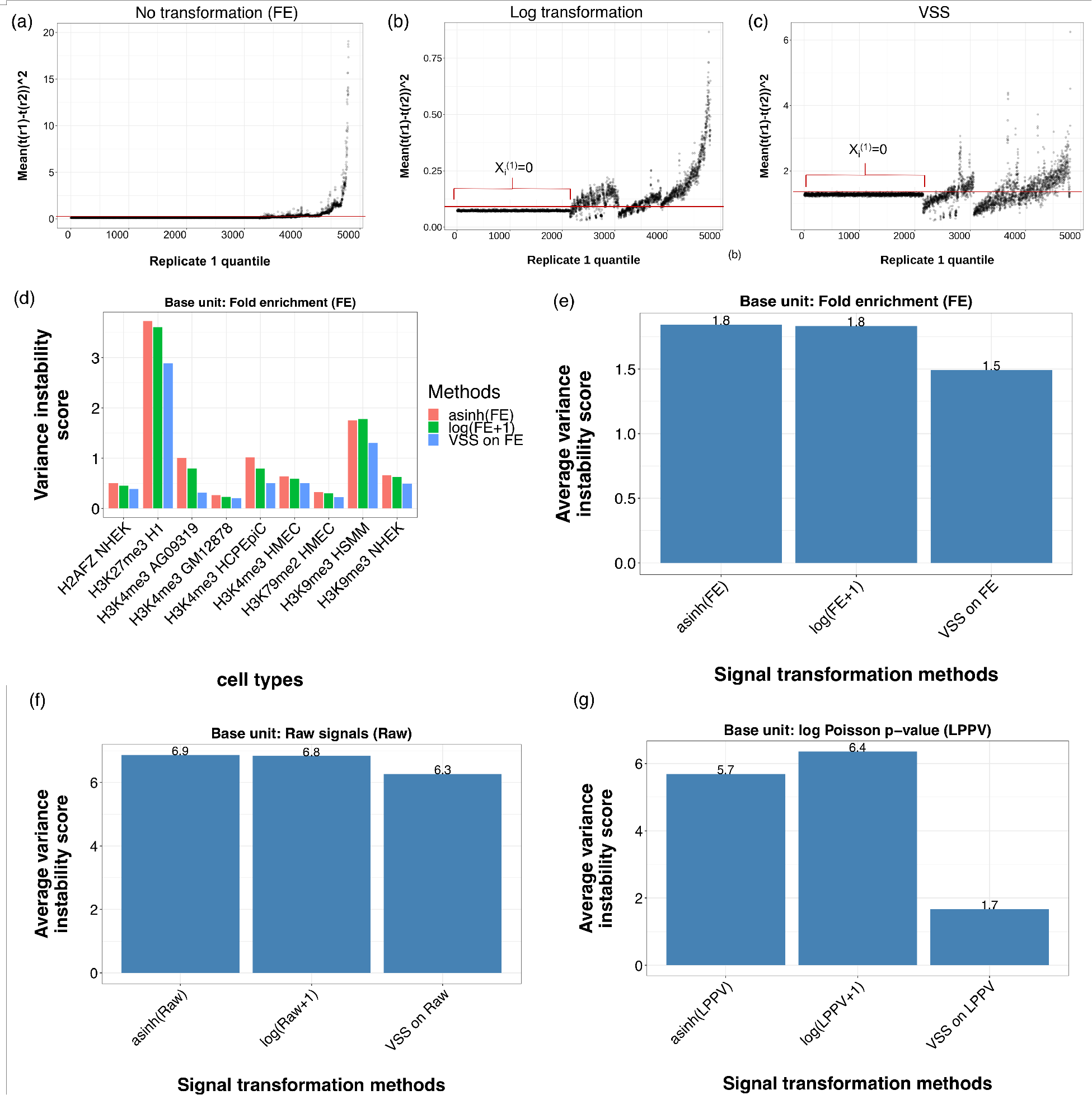
Variance instability of transformed signals. (a,b,c) Each point corresponds to a bin, where binning is defined according to replicate 1 value (Methods). Horizontal axis indicates binning index. Vertical axis indicates squared difference of values between replicates in H3K4me3 HMEC. The flat line on the left half of each plot corresponds to positions where *x*^(1)^ = 0. Signals with stable variance show a flat (constant) trend on this plot; a trend (increasing or decreasing) indicates unstable variance. (a) Untransformed values (Fold enrichment signals). (b) log(*FE* + 1) transform. (c) VSS on FE transformation. (d) Variance instability score on Fold enrichment signals (Methods). Lower values indicate more stable variance. (e) Same as (d), but averaged across experiments. (f) Averaged variance instability score on Raw signals. (g) Averaged variance instability score on log Poisson p-value (LPPV). VSS’s meanvariance relationship was trained on chromosome 22 and variance instability score was computed on chromosome 21. We omitted the variance instability value for untransformed signals because its value would distort the vertical axis (mean of 601, 117 and 1877 across experiments for Fold enrichment signals, raw signals and log Poisson p-value signals respectievly).

### 3.3 VSS signals improve segmentation and genome annotation (SAGA) algorithms

To evaluate the efficacy of transformed signals as input to Gaussian models, we use segmentation and genome annotation (SAGA) as an example. Segmentation and genome annotation algorithms are widely used to integrate genomic data sets and annotate genomic regulatory elements [12–16]. Following previous work [15, 39], we evaluated the quality of an annotation by the correlation between the label of a gene body and whether that gene is expressed as measured by RNA-seq (Methods). We evaluated this correlation across multiple cell types and model initializations (Methods). We believe that high quality input signals will lead to a high quality annotation. We used the SAGA algorithm Segway [12] annotation for this analysis.

We found that using variance-stabilized signals from VSS improves annotations produced by SAGA algorithms (Figure, 4 and 5). As had been previously observed [12], using non-stabilized fold enrichment (FE) signal results in poor performance (mean *r*^2^=0.47, Figure 4). To account for this, Segway recommends using an asinh transform; doing so substantially improves performance (mean *r*^2^ =0.57, Figure 4). VSS produces similar results to asinh on FE data (mean *r*^2^=0.57, *p* = 0.28). However, VSS outperforms asinh when using raw or log Poisson p-value (LPPV) as the base signals (*p* = 0.028 and *p* = 0.007 respectively, paired one-sided Wilcoxon rank-sum test). Likewise, VSS outperforms a log transformation for FE and LPPV signals (*p* = 0.023 and *p* = 0.013 respectively). This improvement likely results from the fact that VSS stabilizes variance in all cases, whereas asinh does so only when data sets happen to have a specific mean-variance relationship.

**Fig. 4.**
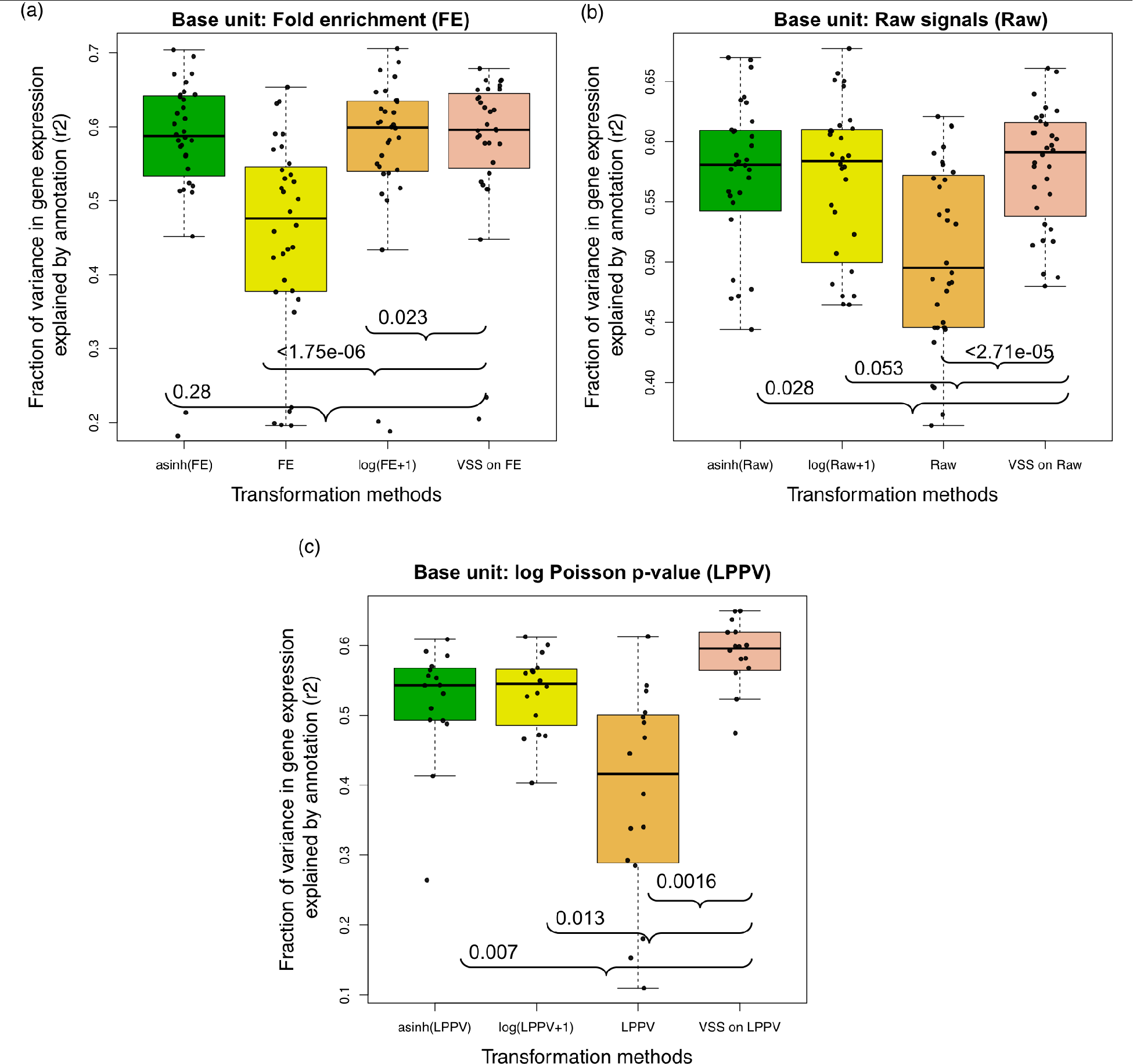
Evaluation of annotations relative to gene expression. Vertical axis is the fraction of variance in gene expression explained (r2, Genome annotation evaluation section). Horizontal axis is the number of features or states in a given model respectively (k). Results are shown on chromosome 21 for an VSS model trained on chromosome 22. Brackets indicate p-value from paired one-sided Wilcoxon rank-sum test.

**Fig. 5.**
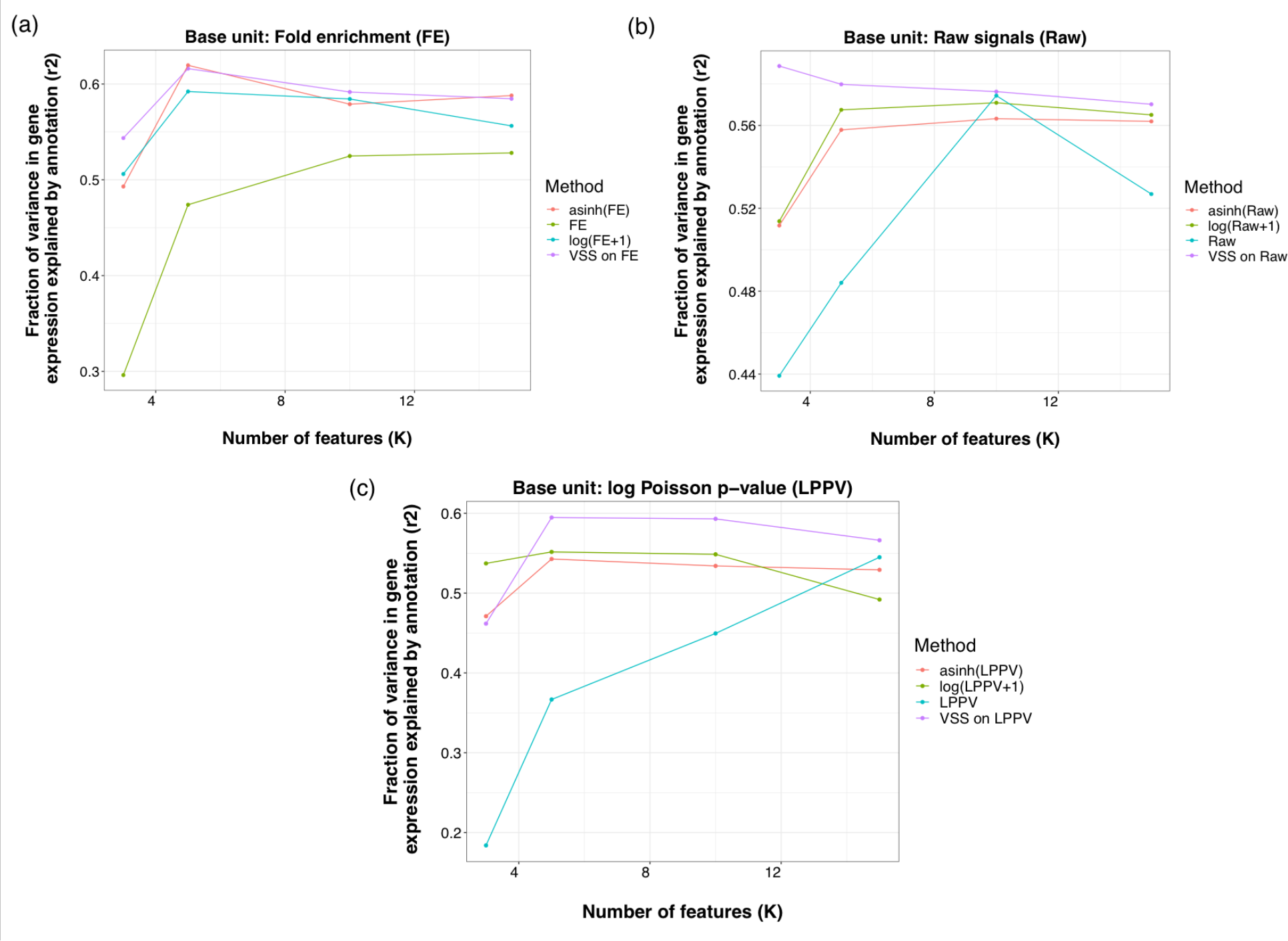
Evaluation of annotations relative to gene expression. Vertical axis is the fraction of variance in gene expression explained (r2, Gene expression evaluation section). Horizontal axis is the number of features or states in a given model respectively (k). Results are shown on chromosome 21 for an VSS model trained on chromosome 22.

## 4 Discussion

In this manuscript we proposed VSS, a method that produces units for sequencing-based genomic signals that have the desirable property of variance stability. We found that the transformations that are currently used to stabilize variance—log(*x* + 1) and asinh(*x*)—do not fully do so. In fact, we found that the mean-variance relationship of genomic signals varies greatly between data sets, indicating that no single transformation can be applied to all data sets uniformly. Instead, variance stability requires a method such as VSS that empirically determines the experiment-specific mean-variance relationship.

We showed that VSS successfully stabilizes variance in genomic data sets. Further, we found that using variance-stabilized data improves the performance of Gaussian models such as SAGA.

Variance-stabilized signals will aid in all downstream applications of genomic signals. In particular, they are valuable for two reasons. First, VSS signals allow downstream methods to use squared error loss or Gaussian likelihood distributions, which are much easier to optimize than the existing practice of implementing a model that accounts for the mean-variance relationship. This will improve tasks that currently use Gaussian models, such as chromatin state annotation and imputation.

Second, VSS signals can be easily analyzed by eye because the viewer does not need to take the meanvariance relationship into account when visually inspecting the data. For example, when viewing genomic signals in a genome browser, variance-unstable signals often exhibit high peaks that swamp the vertical axis and flatten other variations in signal (Figure 6). Existing methods for handling this problem—using a log/asinh transform or cutting off the vertical axis—can also be effective, but they lack the principled basis of VSS.

**Fig. 6.**
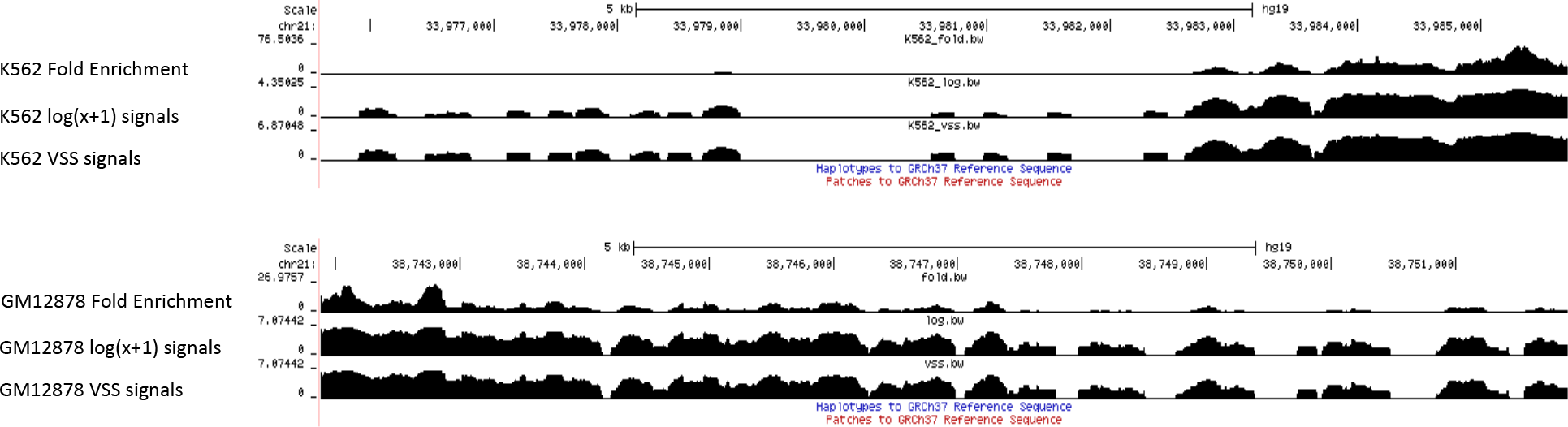
Visualization of genomic signals (H3K4me3) in the UCSC genomic browser.

A key limitation of VSS is that it requires the availability of replicated data. A fruitful direction for future work might aim to remove this dependence, for example by training a consensus transformation to apply across non-replicated data. A related limitation is that VSS relies heavily on the comparability of its input replicates. If there are (e.g.) batch effects between the replicates, VSS may over-estimate variance and produce a poor mean-variance fit.

