## Supplementary information for "VSS: Variance-stabilized signals for sequencing-based genomic signals"

### A Datasets Information

**Table 1.** Genomic assays ENCODE accession numbers.

| Assay/Celltype | ENCODE accession number |
|---|---|
| H3K4me3 GM1287 | ENCSR000AKA |
| H3K4me3 H1-hESC | ENCSR000AMG |
| H3K4me3 HUVEC | ENCSR000AKN |
| H3K4me3 K562 | ENCSR000AKU |
| H3K4me3 NHLF | ENCSR000DWZ |
| H3K4me3 GM06990 | ENCSR000DQV |
| H3K4me3 HCPEpiC | ENCSR000DTN |
| H3K4me3 AG09319 | ENCSR000DPU |
| H3K4me3 NHEK | ENCSR000ALO |
| H3K4me3 HMEC | ENCSR016JWS |
| H3K4me3 HSMM | ENCSR000ANK |
| H3K36me3 H1-hESC | ENCSR925LJZ |
| H3K4me1 H1-hESC | ENCSR631RJR |
| H3K27me3 H1-hESC | ENCSR216OGD |
| H3K9me3 H1-hESC | ENCSR883AQJ |
| H2AFZ NHEK | ENCSR000ARL |
| H2AFZ HSMM | ENCSR000APA |
| H3K79me2 NHEK | ENCSR000ARM |
| H3K79me2 HSMM | ENCSR000ANQ |
| H3K79me2 HMEC | ENCSR000ASB |
| H3K9me3 NHEK | ENCSR000ARN |
| H3K9me3 AG04450 | ENCSR000DPJ |
| H3K9me3 HMEC | ENCSR000ARG |
| H3K9me3 HSMM | ENCSR000ANR |
| H3K36me3 HMEC | ENCSR000ALY |

### B Setting VSS hyperparameters

In order to identify optimum values for VSS's hyperparameters, we evaluated many possible combinations of values using the two evaluation measures, likelihood analysis and variance instability (Figure 7). The set of parameters is considered optimized if they minimize the two evaluation metrics. Thus, based on these results, we chose $\beta = 10^3$ and $b = 10^5$ as the optimal set of the parameters as it satisfies all evaluation metrics simultaneously. We have also compared the optimization results with $\log(x+1)$ transformation which is shown by red lines in the Figure 7. The results indicate that our approach is outperforming the $\log(x+1)$ transformation in all investigated evaluation metrics. The parameter setting results of the first mode is shown in Figure 7. Based on the experiments, we chose to use $\beta = 10^3$ and $b = 10^5$ as the best combination of the parameters for the first mode since it optimizes all evaluation metrics simultaneously.

For the second mode of the mean-variance relationship identification, we considered all the genomic signals in the smoothing procedure rather than considering the fluctuated signals in one bin. Thus, based on two criteria, we chose $\beta = 10^7$ and $b = 10^3$ as the optimal set of the parameters for the second mode as it satisfies all evaluation metrics simultaneously. We have also compared the optimization results with $\log(x+1)$ transformation which is shown by red lines in the Figure 8.

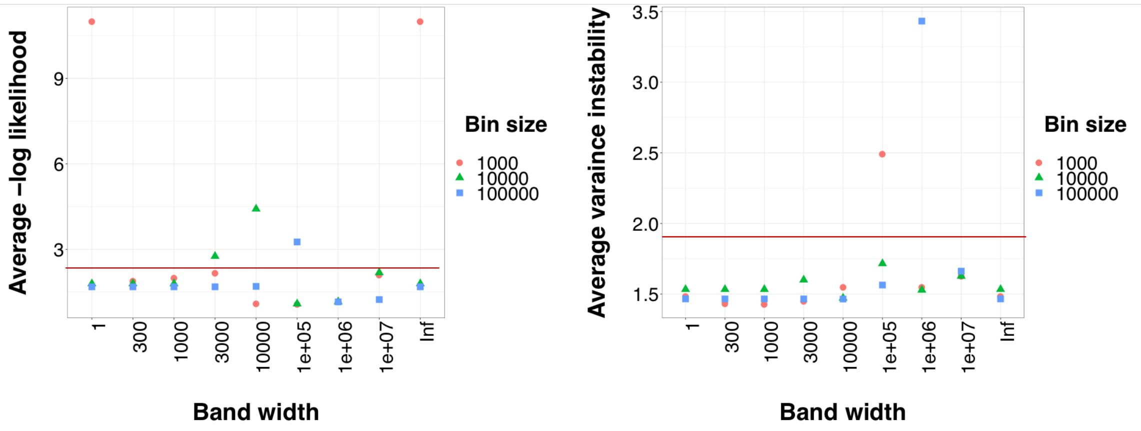


**Fig. 7.** Relationship between bin size and bandwidth in (a) likelihood analysis (b) variance instability analysis. Among the band width values, the value Inf indicates that we applied no smoothing on the curve (Unweighted mean-variance curve). Red line indicates the performance of $\log(x+1)$ transformation.

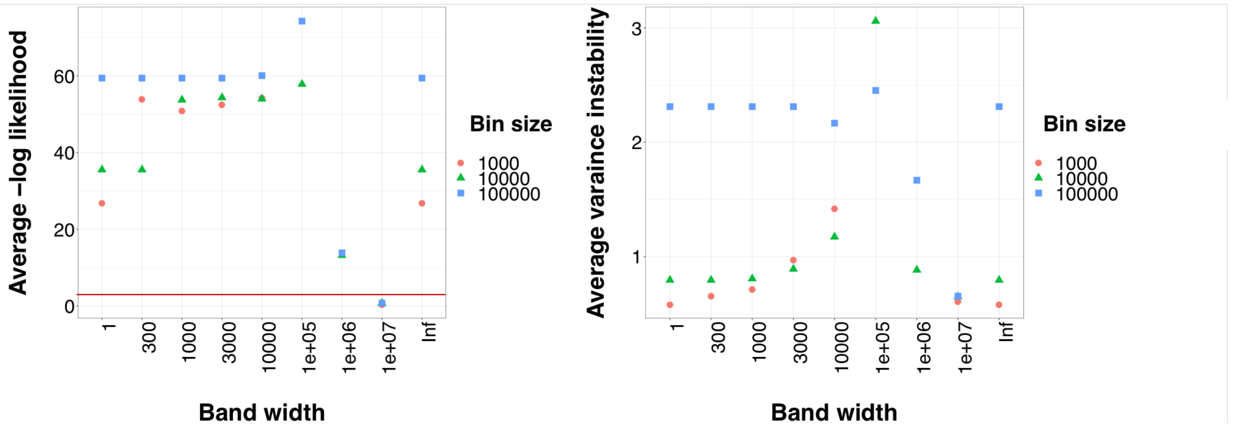


**Fig. 8.** Relationship between bin size and bandwidth in (a) likelihood analysis (b) variance instability analysis. Among the band width values, the value Inf indicates that we applied no smoothing on the curve (Unweighted mean-variance curve). Red line indicates the performance of $\log(x+1)$ transformation. Average variance instability score for $\log(x+1)$ is 6.4 .

#### C Alternative offsets for a log transformation, $\log(ax + b)$

We evaluated whether it is possible to improve the log transform using a linear transform $\log(ax+b)$. We did so using the previously-described log likelihood and variance instability evaluations. We found that no single set of parameters performed best across all data sets (Supplementary Figures 9, 10), and all performed less well than VSS.

we could not identify a single pair of parameters that can optimize both criteria. Therefore, we believe that default choice of $a = 1$ and $b = 1$ can be applied to the $\log(ax + b)$

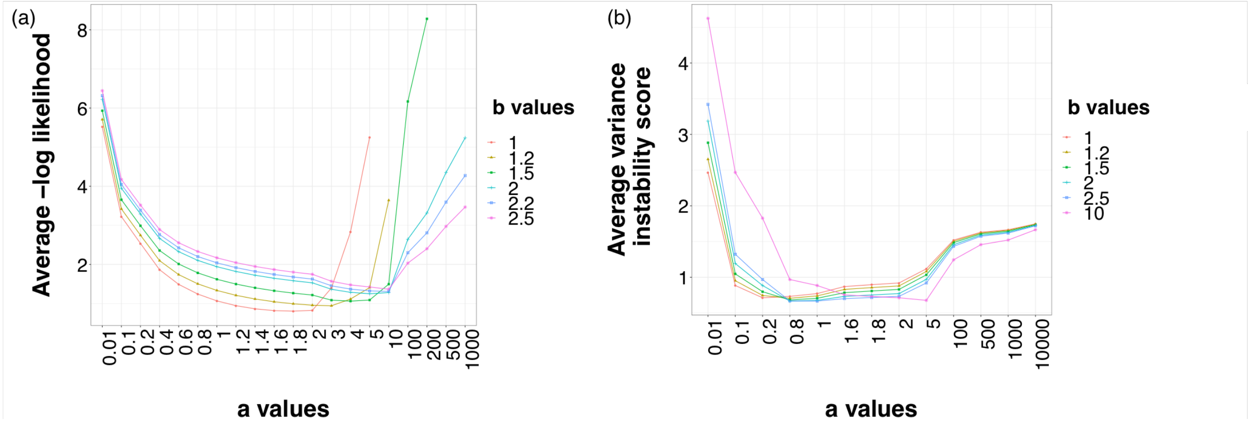


**Fig. 9.** Relationship between $a$ and $b$ in $\log(ax+b)$ transformation in (a) likelihood analysis (b) variance instability analysis .

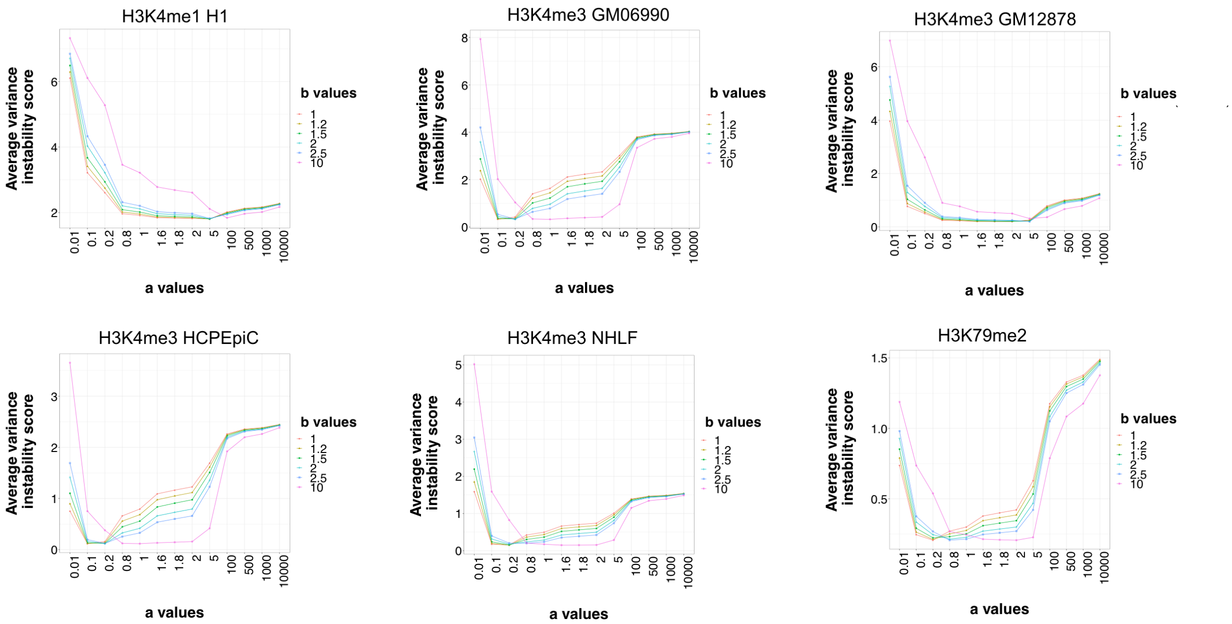


**Fig. 10.** Relationship between $a$ and $b$ in $\log(ax+b)$ transformation in variance instability analysis .
